# Gene regulatory changes underlie developmental plasticity in respiration and aerobic performance in highland deer mice

**DOI:** 10.1101/2022.09.24.509328

**Authors:** Rena M. Schweizer, Catherine M. Ivy, Chandrasekhar Natarajan, Graham R. Scott, Jay F. Storz, Zachary A. Cheviron

## Abstract

Phenotypic plasticity can play an important role in the ability of animals to tolerate environmental stress, but the nature and magnitude of plastic responses are often specific to the developmental timing of exposure. Here, we examine changes in gene expression in the diaphragm of highland deer mice (*Peromyscus maniculatus*) in response to hypoxia exposure at different stages of development. In highland deer mice, developmental plasticity in diaphragm function may mediate changes in several respiratory traits that influence aerobic metabolism and performance under hypoxia. We generated RNAseq data from diaphragm tissue of adult deer mice exposed to 1) life-long hypoxia (before conception to adulthood), 2) post-natal hypoxia (birth to adulthood), 3) adult hypoxia (6-8 weeks only during adulthood), or 4) normoxia. We found five suites of co-regulated genes that are differentially expressed in response to hypoxia, but the patterns of differential expression depend on the developmental timing of exposure. We also identified four transcriptional modules that are associated with important respiratory traits. Many of the genes in these transcriptional modules bear signatures of altitude-related selection, providing an indirect line of evidence that observed changes in gene expression may be adaptive in hypoxic environments. Our results demonstrate the importance of developmental stage in determining the phenotypic response to environmental stressors.

## Introduction

Plasticity in whole-animal physiological performance potentially stems from a combination of environmentally induced changes during pre- and/or postnatal development and adulthood (acclimatization). The nature of plastic responses to particular environmental stimuli are often sensitive to the developmental timing of exposure, with early exposures generally exerting a disproportionate influence on performance phenotypes in adulthood (Hammond et al., 2006). Specifically, environmental perturbations early in development may exert strong influences on the morphological development of organ systems that support whole-organism performance. Once these developmental windows close, the nature and amplitude of plastic responses may be more restricted. As a result of the reduced “degrees of freedom” available for plastic responses, exposure to environmental stressors during later developmental stages may be expected to produce more modest changes in the expression of performance-related traits and may be mediated by different physiological mechanisms (Hammond et al., 2006).

The effects of hypobaric hypoxia on aerobic performance offer a promising system to study life-stage specific impacts of environmental stressors. Hypoxia exposure during development is known to have profound effects on aerobic metabolism and performance by altering physiological traits that govern oxygen uptake, transport, and utilization. Yet, these effects are contingent on the timing of exposure, and are especially pronounced for traits that influence respiratory physiology. For example, the hypoxic ventilatory response (HVR) is often augmented by pre-natal hypoxia exposure but is often reduced by post-natal exposure (Gleed & Mortola, 1991; Ivy & Scott, 2021; Peyronnet et al., 2000, 2007). These life stage-specific responses may stem from perturbation of critical developmental windows for key components of the hypoxic chemoreflex, such as the carotid body and neural networks that underlie the control of breathing (Bavis, 2005; Brooks & Tenney, 1968; Lumbroso & Joseph, 2009; Sterni et al., 1999; Wong-Riley et al., 2019; Yilmaz et al., 2005).

Detailed studies of plastic responses to environmental challenges at different life stages can provide key insights into the limits of phenotypic plasticity, and the specific mechanisms that underlie those limits. On one hand, we may expect the same core set of genes to underlie plastic physiological changes across life stages, with earlier developmental exposure to hypoxia eliciting a greater magnitude of expression changes in this core set. Early exposure may allow more time to initiate transcriptomic responses, and continued exposure could reinforce these regulatory changes, allowing for more pronounced regulatory differentiation between normoxia and hypoxia exposed individuals. Under this hypothesis, the set of genes that differentiate hypoxia-exposed individuals from normoxia-exposed individuals would be the same across life stages, and only the magnitude of expression differences of this core set of genes would vary (and would correlate with time since exposure). Alternatively, distinct gene expression phenotypes may distinguish hypoxia-exposed individuals depending on the developmental timing of exposure. This pattern would suggest that either distinct sets of genes underlie plastic responses at different developmental timepoints, or that a common core set of genes is only transiently differentially expressed.

The deer mouse (*Peromyscus maniculatus*) is well-suited for studies exploring phenotypic plasticity and its underlying mechanisms. Deer mice that are native to high elevations in the western US have evolved increased aerobic performance (*V*O_2 max_) under hypoxia compared to lowland conspecifics and closely related species (Chappell & Snyder, 1984; Cheviron et al., 2013; Cheviron et al., 2012; Cheviron et al., 2014; Lui et al., 2015; Tate et al., 2017, 2020). The enhanced aerobic performance under hypoxia is associated with increased survivorship in free-ranging mice at high elevation (Hayes & O’Connor, 1999) and is associated with plastic and genetically based modifications across the oxygen transport cascade (Ivy & Scott, 2015; Schweizer et al., 2021, 2019; Storz & Cheviron, 2020; Storz et al., 2019; Storz & Scott, 2019). Research into the ontogeny of these differences has only just begun, but some components of the unique physiology of highlanders is clearly manifest within the first few weeks of life (Ivy et al., 2020; Ivy et al., 2021; Robertson et al., 2019; Velotta et al., 2020). The plastic responses to hypoxia also differ between highland and lowland mice. For example, hypoxia exposure during adulthood improves ventilation in lowlanders, but it has no effect on the ventilatory phenotypes of highlanders (Ivy & Scott, 2017; Ivy et al., 2021; Ivy & Scott, 2017).

In a recent study of highland deer mice from the Rocky Mountains, Ivy et al. (2021) found that exposure to hypoxia during early development improved both ventilation and whole-animal aerobic performance (cold-induced VO_2_max) during hypoxia in adulthood, and the magnitude of these improvements depended on the stage of developmental exposure. Normoxic controls were compared to mice exposed to hypoxia starting at three different life stages and extending into adulthood: life-long hypoxia (before conception to adulthood); post-natal hypoxia (birth to adulthood); and adult hypoxia (6-8 weeks only during adulthood). Mice exposed to hypoxia during pre-natal development outperformed those exposed to hypoxia starting later in early post-natal life, and hypoxia exposure during both developmental stages improved performance to a greater extent than hypoxia acclimation during adulthood. Developmental exposure to hypoxia also had pronounced effects on breathing phenotypes in highlanders, which likely contributed to the improvements in aerobic performance. Namely, pre- and post-natal exposure increased total ventilation at VO_2_max compared to normoxia-exposed controls by 100% and 63%, respectively, but ventilation was unaffected by hypoxia when adults were acclimated to hypoxia. These improvements in total ventilation were driven by increases in tidal volume, and together contributed to significant improvements in arterial oxygen saturation under hypoxia compared to controls. Interestingly, mice exposed to hypoxia during adulthood also improved arterial oxygen saturation, but this improvement was not attributable to changes in ventilatory phenotypes.

In this study, we examine gene expression changes in the diaphragm that underlie plastic responses to hypoxia exposure at different life stages. We focus on the diaphragm because of its central role in mammalian ventilation, and because our previous studies of highland deer mice suggest that developmental plasticity in diaphragm form and function may mediate changes in several respiratory traits that contribute to aerobic metabolism and performance under hypoxia (Catherine M Ivy et al., 2021). We address three interrelated questions: 1) What are the underlying gene regulatory changes that contribute to plastic responses to hypoxia in early development and adulthood?; 2) How are gene regulatory phenotypes associated with ventilatory traits in response to hypoxia exposure?; and 3) Have genes associated with ventilatory traits at different life stages been subject to positive selection in highland mice? We show that several suites of co-regulated genes are differentially expressed in response to hypoxia, but the patterns of differential expression depend on the developmental timing of exposure. We also identify several transcriptional modules that are associated with ventilatory traits and whole-organism performance, and that many of the genes in these transcriptional modules bear signatures of elevation-related selection. Together these results provide new insights into the genetic and regulatory bases of adaptive developmental plasticity.

## Materials and Methods

### Experimental procedures and phenotyping

The experimental animals were second-generation descendants of wild-caught mice from the summit of Mt. Evans (4350 m), Clear Creek County, Colorado, USA. As previously described (Catherine M Ivy et al., 2021), captive mice were exposed to hypoxia, using custom-designed hypobaric chambers. The experiments involved four treatment groups: 1) Life-long hypoxia (LH) mice were conceived, born, and raised to adulthood in hypobaric hypoxia (~12 kPa); 2) Post-natal hypoxia (PNH) mice were conceived and born in normoxia, then the family moved into hypobaric hypoxia within 12 hours from birth; 3) Adult hypoxia (AH) mice were conceived, born, and raised to adulthood in normoxia (~21 kPa), then acclimated for 6-8 weeks to hypobaric hypoxia; and 4) Control (C) mice were conceived, born, and raised into adulthood in normoxia, and kept at normoxia for the duration of the experiment. We obtained phenotypic data from previously published studies for almost all individuals (Ivy et al., 2021; Ivy & Scott, 2021). Cold-induced VO_2_max (ml O_2_ g^−1^ min^−1^) was measured in mice at both 21 kPa O_2_ and 12 kPa O_2_, along with additional cardiorespiratory measurements at VO_2_max, including arterial oxygen saturation (%), heart rate (beats min^−1^), breathing frequency (breaths min^−1^), tidal volume (μl g^−1^), total ventilation (ml g^−1^ min^−1^), air convection requirement (the quotient of total ventilation and VO_2_max; ml air ml O_2_^−1^), and pulmonary extraction (%). Resting measurements of breathing frequency (breaths min^−1^), tidal volume (μl g^−1^), and total ventilation (ml g^−1^ min^−1^) were also made at room temperature at 21 kPa O_2_, 12 kPa O_2_, and 8 kPa O_2_. Body mass (g) and diaphragm mass (g) were also measured. Further methods for these phenotype data are available in these earlier publications (Ivy et al., 2021; Ivy & Scott, 2021).

### RNA sequencing of diaphragm tissue

We used high-throughput RNA sequencing to identify gene expression differences among individuals exposed to different hypoxia (LH, PNH, AH) or normoxia (C) treatments. For each sample, Azenta Life Technologies (Plainsfield, NJ) extracted RNA from diaphragm tissue using a SMART-Seq HT kit for full-length cDNA synthesis and amplification, then prepared sequencing libraries using an Illumina Nextera X kit. After checking RNA and library quality using a TapeStation, Azenta performed 150 bp paired end sequencing on a HiSeq 4000 machine. Twelve control samples were prepared and sequenced together in one lane, followed by the remaining 27 samples prepared and sequenced on another lane.

First, we removed adapters from sequence reads by using BBduk 38.93 (sourceforge.net/projects/bbmap/) with default parameters other than trimming to the right with ktrim=r’ and a kmer length of 24. Then, we used trimmomatic (default parameters other than SLIDINGWINDOW:5:20) (Bolger, Lohse, & Usadel, 2014) to remove bases below quality 3 at the beginning and end of the read, then further trim bases when the average quality per 5-base window dropped below 20. Subsequent filtered reads were mapped to the chromosome-level assembly of the nuclear and mitochondrial *Peromyscus maniculatus* genomes GCA_003704035.1) using *HiSat2 2.2.1* (Zhang et al., 2021) with a maximum and minimum mismatch penalty (--mp 2,0). We implemented *featureCounts* v1.5.2 (Liao et al., 2014) to count numbers of reads aligning to annotated genes (n=25,431).

We processed and analyzed count data using the R programming language. A record of all commands and output is available within an R markdown file (https://github.com/renaschweizer/pman_diaphragm_rnaseq). To detect any potential technical artifacts or batch effects in our data, we ran a principal components analysis (PCA) using five technical sequencing metrics (i.e., percent GC content, mean quality score, percent bases with quality score greater than or equal to 30, genome alignment rate, percent reads assigned with fatureCounts) and the base *prcomp* function. Using ggbiplot v 0.55 https://github.com/vqv/ggbiplot/tree/experimental), we plotted PC1 and PC2, then identified and removed four samples (N1, N2, AH1, AH2) located outside of the 95% data ellipse. To reduce measurement error caused by genes with low numbers of reads, we removed genes with fewer than 20 average reads across individuals (n= 15,675 genes remaining). We normalized read counts among libraries in the remaining samples, then log-transformed using the *calcNormFactors* and *cpm* functions, respectively, within the *edgeR* package (Robinson et al., 2009).

### Test for treatment effects on gene expression at gene-level

We searched for differential gene expression (i.e., for individual genes) using three contrasts: 1) a “lifelong” contrast of the control vs lifelong hypoxia treatments, 2) a “postnatal” contrast of the control vs. post-natal hypoxia treatments, and 3) an “adult” contrast of the control vs. adult hypoxia treatments. First, we used the function *estimateGLMTagwiseDisp* to estimate gene dispersion using a design matrix with a column for each of the five treatment groups. Then, we used a generalized linear model to test differences in normalized transcript abundance (the response variable), with treatments as factors. We assessed significance using quasi-likelihood F tests (implemented with the *glmQLFit* and *glmQLFTest* functions), and corrected for multiple testing using a genome-wide false discovery rate threshold of q ≤ 0.05 (Benjamini & Hochberg, 1995). We tested for enrichment of gene ontology categories within significantly differentially expressed gene sets using the *gost* function within *gProfiler* (Raudvere et al., 2019). The *Mus musculus* gene ID for each *Peromyscus maniculatus* gene was used for the query, and the background gene list was a custom set of all expressed genes within diaphragm tissue. We used a false-discovery rate threshold of 0.05 to assign significance.

### Test for treatment effects on gene regulatory modules

We predicted that developmental hypoxia exposure is associated with unique expression signatures that differ from those induced by hypoxia exposure during adulthood. To test for treatment effects on gene expression, we implemented a weighted gene co-expression network analysis (WGCNA) approach within the *WGCNA* package in R (Langfelder & Horvath, 2008). Briefly, WGCNA identifies clusters of genes that have correlated expression profiles (called co-expression modules). We used the *blockwiseModules* function in WGCNA to define modules, using a maximum block size of 16000 (greater than the total number of genes) and setting the random seed for reproducibility. We used a soft thresholding power of 13, since this was the initial inflection point at which the scale free topology model fit began to decrease with higher thresholding powers (Figure S1). Co-expression modules were ultimately defined using a hierarchical clustering approach, as in Velotta et al. (2020). For all co-expression modules defined through the WGCNA algorithm, we tested for a significant effect of treatment on expression using an ANOVA of a linear mixed-effects model with eigengene value as the response variable, treatment group as a fixed-effect variable, and family as a random-effect variable. The eigengene, which is the first principal component of a module, is symbolic of the gene expression data for a module. We explored group differences among modules that showed treatment effects using post-hoc Tukey tests. Statistical analyses were performed using the *lmer* function in the *lmerTest* package in R (Kuznetsova et al., 2017), the *anova* function in base R, and the *emmeans* function in the *emmeans* package (https://cran.r-project.org/web/packages/emmeans/index.html). Because highly connected genes may have particular functional importance in regulating trait expression, we identified the top hub gene in each of our significant modules using the chooseTopHubInEachModule function in WGCNA. As with the DE gene set, we tested for enrichment of gene ontology categories within modules with significant treatment effects using gene ontology enrichment analysis.

### Correlation of gene regulatory modules with phenotypes

We measured the Pearson correlation between the module eigengene values from WGCNA and the phenotypic trait data using the *cor* function of WGCNA. We assigned p-values using the *corPvalueStudent* function, then corrected for multiple-testing using the p.*adjust* function and the Benjamini & Hochberg correction (1995). We used a false-discovery rate threshold of 0.05 to assign significance.

### Test for overlap with genes under positive selection at high altitude

In a genome-wide survey of DNA polymorphism that included the same high-elevation source population from which our experimental mice were derived, we previously documented evidence for positive directional selection in a set of 1852 genes (Table S1) (Schweizer et al., 2021). These candidate genes for high-elevation adaptation were identified by calculating the Population Branch Statistic (PBS; Yi et al., 2010) in exome-wide windows of 5 kb, followed by the use of a demographic model to identify outlier loci at a significance threshold of 99.9%. We examined whether this set of selection-nominated candidate genes included any of the genes in our treatment-significant WGCNA modules or those in our set of significantly differentially expressed genes.

## Results

### Differential gene expression across early development and adulthood - edgeR

We obtained RNAseq data for 35 samples (n_AH_=8; _nLH_=8; n_N_=10; n_pNH_=9), with a mean genome alignment rate of 85.39% ± 2.39% and a mean FeatureCounts assignment rate of 58.33% ± 13.10% (see additional quality metrics in Table S2). Using EdgeR, we identified hundreds to thousands of significantly differentially expressed (DE) genes across treatment groups relative to the control samples (Table 1; Table S3). Only a single gene was exclusive to the lifelong hypoxia group, and overall, the lifelong hypoxia group showed a smaller number of DE genes (n=222 at FDR≤0.05) than the post-natal group (n=2919 genes at FDR≤0.05) and the adult hypoxia group (n=6409 genes at FDR≤0.05) (Figure 1). 203 genes were exclusively identified in the PNH group, and 209 genes were exclusive to the developmental groups. All of the 207 genes that overlapped amongst all three comparisons showed a consistent direction of expression relative to the control groups (Figure 2).

**Table 1.**
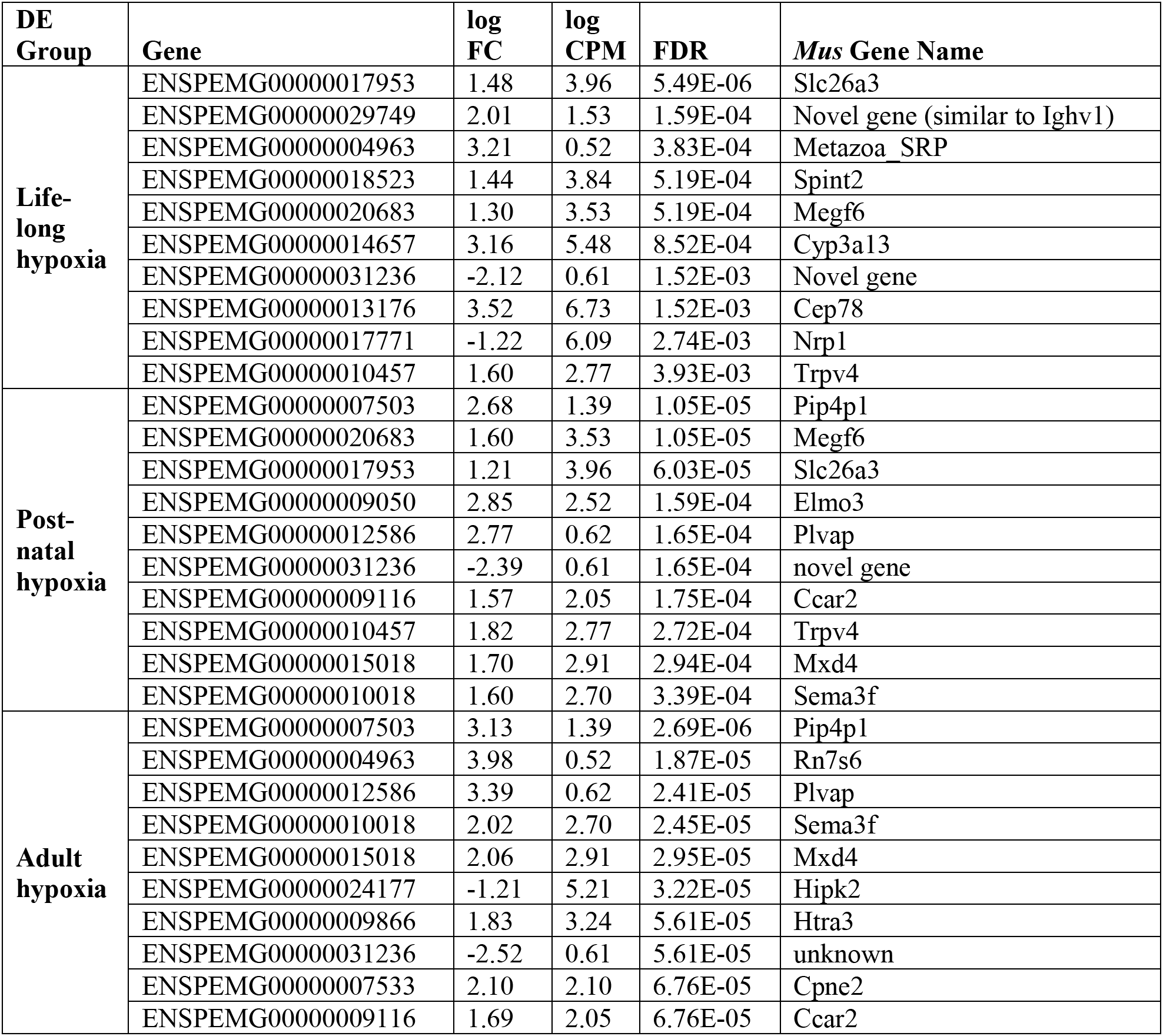
Top ten differentially expressed genes for life-long hypoxia, post-natal hypoxia, and adult hypoxia relative to control. For each gene, the Ensembl ID (Gene), log FC (log 2 fold change relative to control), log CPM (log 2 counts per million), FDR (False Discovery Rate-adjusted p-value), and *Mus* gene name is provided.

| DE Group | Gene | log FC | log CPM | FDR | *Mus* Gene Name |
| --- | --- | --- | --- | --- | --- |
| Life-long hypoxia | ENSPEMG00000017953 | 1.48 | 3.96 | 5.49E-06 | Slc26a3 |
|  | ENSPEMG00000029749 | 2.01 | 1.53 | 1.59E-04 | Novel gene (similar to Ighv1) |
|  | ENSPEMG00000004963 | 3.21 | 0.52 | 3.83E-04 | Metazoa SRP |
|  | ENSPEMG00000018523 | 1.44 | 3.84 | 5.19E-04 | Spint2 |
|  | ENSPEMG00000020683 | 1.30 | 3.53 | 5.19E-04 | Megf6 |
|  | ENSPEMG00000014657 | 3.16 | 5.48 | 8.52E-04 | Cyp3a13 |
|  | ENSPEMG00000031236 | -2.12 | 0.61 | 1.52E-03 | Novel gene |
|  | ENSPEMG00000013176 | 3.52 | 6.73 | 1.52E-03 | Cep78 |
|  | ENSPEMG00000017771 | -1.22 | 6.09 | 2.74E-03 | Nrp1 |
|  | ENSPEMG00000010457 | 1.60 | 2.77 | 3.93E-03 | Trpv4 |
| Post-natal hypoxia | ENSPEMG00000007503 | 2.68 | 1.39 | 1.05E-05 | Pip4p1 |
|  | ENSPEMG00000020683 | 1.60 | 3.53 | 1.05E-05 | Megf6 |
|  | ENSPEMG00000017953 | 1.21 | 3.96 | 6.03E-05 | Slc26a3 |
|  | ENSPEMG00000009050 | 2.85 | 2.52 | 1.59E-04 | Elmo3 |
|  | ENSPEMG00000012586 | 2.77 | 0.62 | 1.65E-04 | Plvap |
|  | ENSPEMG00000031236 | -2.39 | 0.61 | 1.65E-04 | novel gene |
|  | ENSPEMG00000009116 | 1.57 | 2.05 | 1.75E-04 | Ccar2 |
|  | ENSPEMG00000010457 | 1.82 | 2.77 | 2.72E-04 | Trpv4 |
|  | ENSPEMG00000015018 | 1.70 | 2.91 | 2.94E-04 | Mxd4 |
|  | ENSPEMG00000010018 | 1.60 | 2.70 | 3.39E-04 | Sema3f |
| Adult hypoxia | ENSPEMG00000007503 | 3.13 | 1.39 | 2.69E-06 | Pip4p1 |
|  | ENSPEMG00000004963 | 3.98 | 0.52 | 1.87E-05 | Rn7s6 |
|  | ENSPEMG00000012586 | 3.39 | 0.62 | 2.41E-05 | Plvap |
|  | ENSPEMG00000010018 | 2.02 | 2.70 | 2.45E-05 | Sema3f |
|  | ENSPEMG00000015018 | 2.06 | 2.91 | 2.95E-05 | Mxd4 |
|  | ENSPEMG00000024177 | -1.21 | 5.21 | 3.22E-05 | Hipk2 |
|  | ENSPEMG00000009866 | 1.83 | 3.24 | 5.61E-05 | Htra3 |
|  | ENSPEMG00000031236 | -2.52 | 0.61 | 5.61E-05 | unknown |
|  | ENSPEMG00000007533 | 2.10 | 2.10 | 6.76E-05 | Cpne2 |
|  | ENSPEMG00000009116 | 1.69 | 2.05 | 6.76E-05 | Ccar2 |

**Figure 1.**
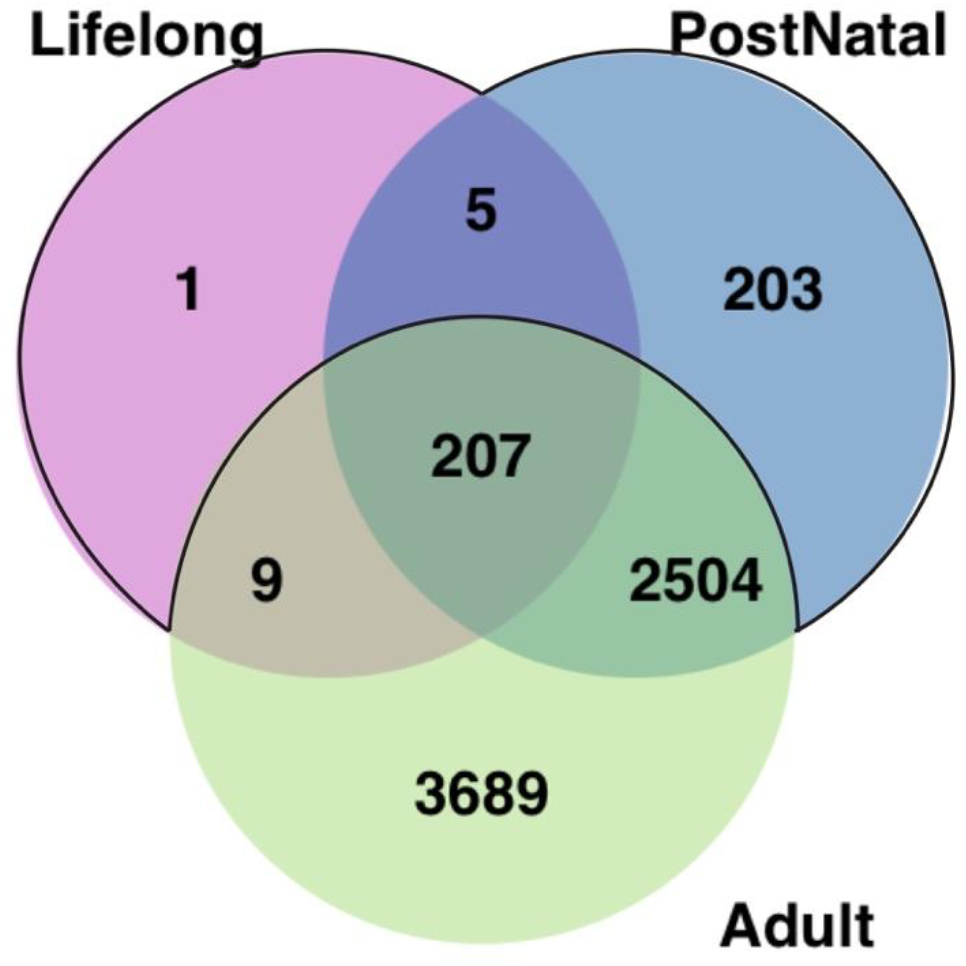
Intersection of significantly differentially expressed genes for each treatment group relative to the control group. Black outline indicates genes that are expressed during early development only (n=209).

**Figure 2.**
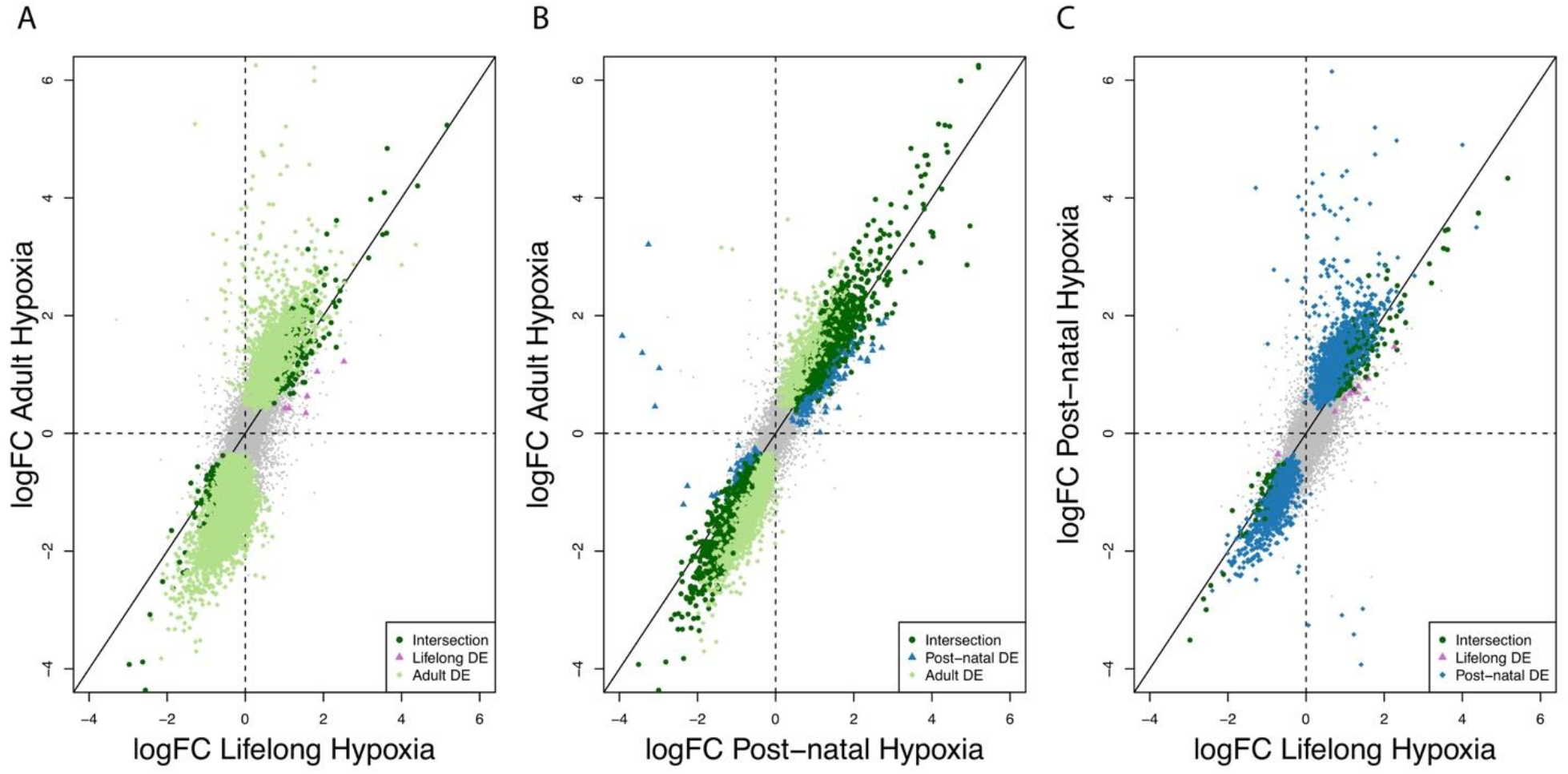
Concordant log fold-change (logFC) amongst 15675 genes (gray points) across three pairwise comparisons (A: Lifelong vs. Adult; B: Post-natal vs. Adult; C: Lifelong vs. Post-natal). For each plot, genes that are significantly differentially expressed relative to control treatment are colored according to whether they are DE in one or the other treatment, or both (see keys). The solid line represents y=x, indicating the same expression in each treatment relative to control.

Although the lifelong hypoxia group had 222 DE genes, there were no significantly enriched GO categories, and three Reactome categories related to immune response and “cell surface interactions at the vascular wall” (Table S4). Seventy-one GO categories that were enriched in the post-natal hypoxia group (n=2919 genes) included processes related to “biological regulation”, “anatomical structure morphogenesis,” “cellular response to oxygen-containing compound”, and immunity/MHC categories. In the adult hypoxia group (n=6409), 41 GO categories were enriched, including several related to “signaling” and “sensory perception,” as well as olfaction (Table S4). There was one significant GO category (“localization”) enriched in the set of intersection genes (n=207), while genes that were exclusively post-natal (n=203) were related to immune function and signaling (Table S4). Finally, DE genes that are exclusive to the post-natal and lifelong hypoxia groups (n=209) showed enrichment of categories related to immune response, major histocompatibility complex, and cell killing.

### Correlated transcriptional expression modules are affected by hypoxia treatment

We identified regulatory mechanisms that potentially underlie responses to hypoxia by identifying coregulated sets of genes (modules in WGCNA) with significant treatment effects. Specifically, WGCNA clustered 13,824 genes into five modules (Table 2); 1851 genes were not assigned to a module. Three of the modules (M1, M2, M3) together contained 83.4% of all genes in the expressed transcriptome. The M1, M2, M3, and M4 modules also showed a significant effect by treatment (Table 2). The M1 module was differentially expressed between mice in the control and post-natal and adult hypoxia groups (C vs. PNH, p-value: 0.0307; C vs. AH, p-value: 0.0011, post-hoc Tukey test). This module was also differentially expressed between the lifelong and adult hypoxia treatments (LH vs. AH, p-value: 0.0107; Figure 3). The M2 module was differentially expressed between the control and adult hypoxia groups (AH vs. C, p-value: 0.0341). Likewise, the M3 module was differentially expressed between the control and adult hypoxia groups (C vs. AH, p-value: 0.0237), and between the control and post-natal hypoxia mice (C vs. PNH, p-value: 0.0345).

**Table 2.** Results of ANOVA for linear mixed effect models of module eigengene value ~ treatment with mouse family ID as a random effect. DenDF is the denominator degrees of freedom calculated by the Satterthwaite method. The numerator degrees of freedom was 3 for all models. Bold *P*-values indicate those values below a significance of 0.05.

| Module | Number of Genes | DenDF | *F*-value | *P*-value |
| --- | --- | --- | --- | --- |
| M1 | 5873 | 27.299 | 8.003 | **0.00055** |
| M2 | 6033 | 27.43 | 4.13 | **0.01544** |
| M3 | 1255 | 29 | 4.571 | **0.00969** |
| M4 | 561 | 27.571 | 3.84 | **0.0204** |
| M5 | 102 | 28.066 | 1.023 | 0.3973 |

**Figure 3.**
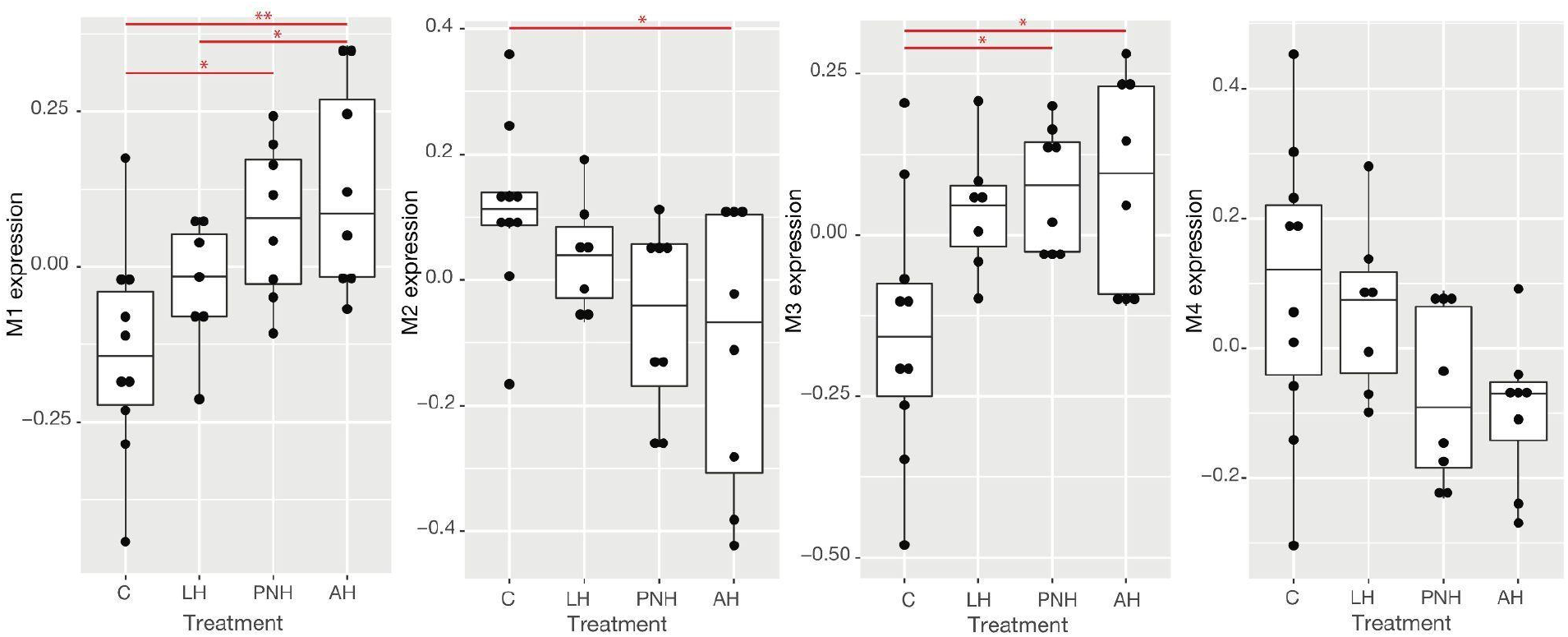
WGCNA module expression by treatment group. Red bars indicate significant difference of means as determined by a post-hoc Tukey test (*: p-value<0.05; **: p-value<0.001).

### Association of correlated transcriptional expression modules with diaphragm mass, arterial O2 saturation, and ventilatory phenotypes

We also aimed to identify gene expression modules that were associated with phenotypes related to respiratory physiology (Table S5). Eigengene values of three modules (M1, M2, M3) were correlated with the arterial O_2_ saturation (M1: r=0.45,p-value: 0.0094; M2: r=−0.42,p-value: 0.017; M3: r=0.46,p-value: 0.0074) (Figure 4). Module M1 was also positively correlated with tidal volume (r=0.36; p-value: 0.04) and total ventilation (r=0.4; p-value: 0.02) of animals at rest at 21 kPa, whereas module M2 was negatively correlated with the latter trait (r=−0.35; p-value: 0.05) (Figure 4). Module M4 was negatively correlated with relative diaphragm mass (r=–0.38; p-value: 0.03) (Figure 4), as well as tidal volume (r=–0.37; p-value: 0.03) and total ventilation (r=−0.35; p-value: 0.05) of animals at rest at 12 kPa. The remaining module (M5) was not significantly correlated with any phenotype, and was not significantly affected by treatment.

**Figure 4.**
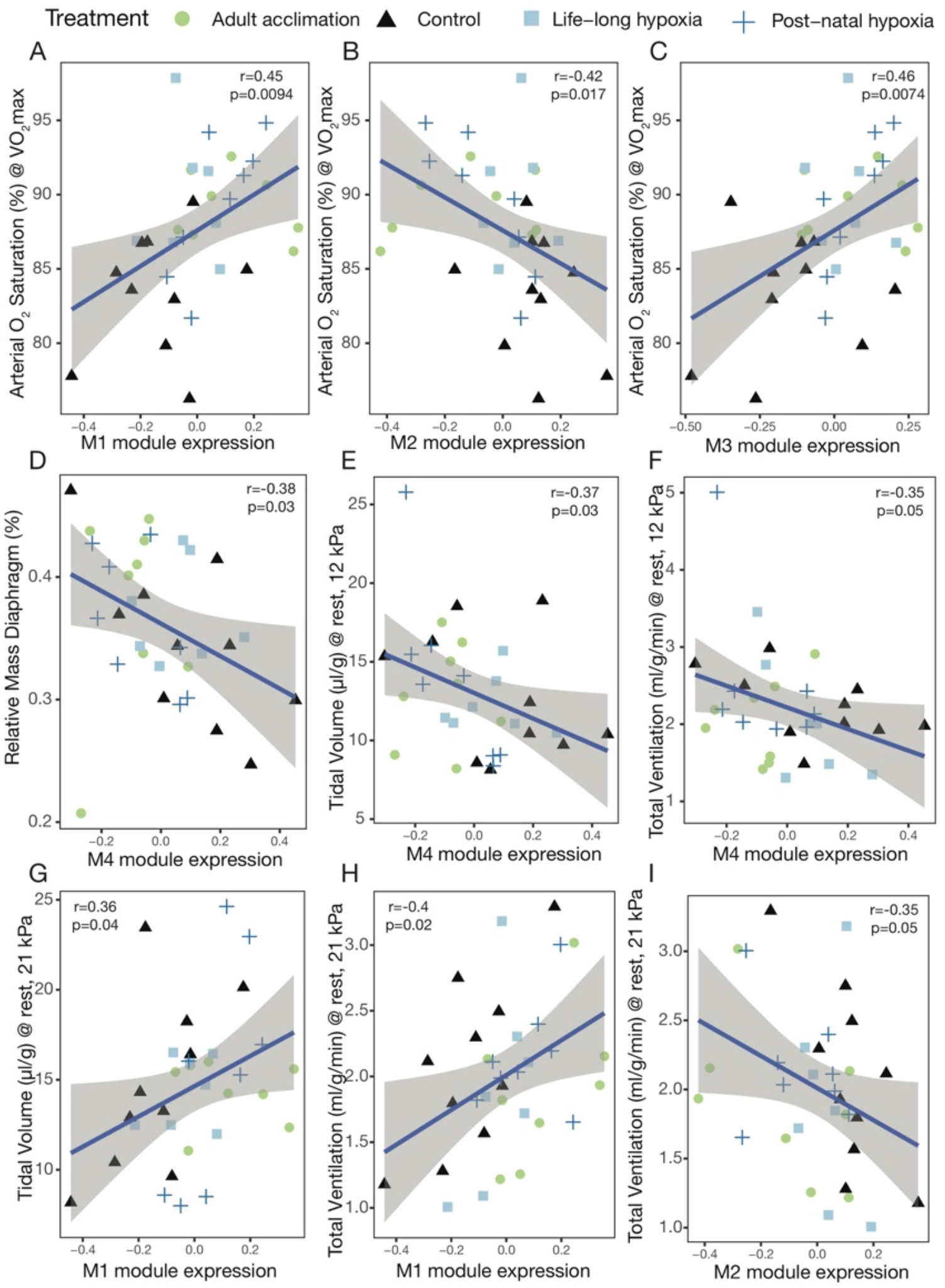
Correlations between physiological phenotypes and expression of diaphragm-specific transcriptional modules. Arterial O_2_ saturation is correlated with expression of modules M1, M2, and M3 (A,B,C) and relative mass of diaphragm (calculated as the mass of the diaphragm divided by the mass of the animal) is correlated with expression of module M4. (D). Tidal volume (E) and total ventilation (F) measured at rest at 12 kPa is correlated with M4 expression, while tidal volume (G) and total ventilation (H) measured at rest at 21 kPa is correlated with M1 module expression. Total ventilation at rest at 21kPa is also correlated with M2 expression (I). The gray shaded area in each panel shows the 95% confidence interval for the linear regression.

We used gene ontology enrichment analysis to assign putative functional categories to genes contained within each of the four modules showing a significant association with a phenotype and a significant effect of treatment on expression (Table S6). The M2 module was enriched for categories related to metabolic processes, such as “mitochondrion organization,” “aerobic respiration,” “oxidative phosphorylation,” and “thermogenesis”. In the M1 module, categories that were enriched were related to sensory perception, reproduction, and vasodilation (e.g., “ion transport,” “sensory perception of chemical stimulus,” “arachidonic acid epoxygenase/monooxygenase process”). The M3 module was enriched for categories related to regulation of gene expression and metabolic processes Finally, in the M4 module, enriched GO categories included those related to gene expression and its regulation as well as regulation of metabolic processes. Detailed information on significantly enriched GO categories and genes included within is provided in Table S6).

Using the chooseTopHubInEachModule function in WGCNA, we identified the top hub gene in each of our significant modules. Clstn2 (Calsyntenin 2), Rex1bd (Required For Excision 1-B Domain Containing), Ccnl2 (Cyclin L2), and Zfp609 (Zinc Finger Protein 609) were the genes with the most connections in the M1, M2, M4, and M3 modules, respectively.

### Trait-correlated gene expression modules contain selection candidates

We found numerous genes within our set of differentially expressed genes that were previously identified as candidates for high-altitude adaptation in deer mice (Schweizer et al., 2021). Sixteen candidate genes overlapped with the “developmental only” set of DE genes (Table S7), and 23 candidate genes overlapped with the three-way intersection set of DE genes (Table S8). Within our four significant WGCNA modules, we found that 662, 407, 150, and 63 genes from the list of outliers overlapped with the M1, M2, M3, and M4 modules, respectively, (Table S9) none of which represented a significant positive enrichment (results not shown). We did find multiple enriched GO categories (Table S10).

## Discussion

Aerobic performance is severely constrained under environmental hypoxia, but these performance decrements can be at least partially mitigated by both developmental plasticity and acclimatization during adulthood. In this study, we exposed deer mice to hypoxia at different developmental stages to determine the gene regulatory changes underlying the outcome of these exposures on performance-related traits in adulthood. Our study had three primary objectives. First, we assessed whether hypoxia exposure at different life stages produced distinct gene expression signatures in adulthood. Second, we identified gene regulatory phenotypes that are associated with respiratory traits. Third, we tested whether genes whose expression was associated with respiratory physiology in hypoxia have experienced a history of positive selection at high elevation.

### What are the underlying gene regulatory changes that contribute to plastic responses to hypoxia in different life stages?

While a core set of 207 genes responded to hypoxia consistently across all experimental exposures, the vast majority of differentially expressed genes were unique to exposure in a particular developmental period (Figure 1, Figure 2). Moreover, the total number of differentially expressed genes varied considerably among the hypoxia exposures, with the greatest gene expression changes occurring in mice exposed to hypoxia during adulthood. Mice exposed to hypoxia throughout life (LH group) exhibited gene expression profiles that were most similar to the control group. A similar pattern was observed for the 207 genes that responded consistently across hypoxia exposures. The comparison between adult hypoxia and control groups showed the greatest difference in expression (mean log FC = 1.56), whereas the comparison between the lifelong hypoxia and control groups showed the smallest difference (mean log FC = 1.39).

These results are not consistent with the idea that persistent differential expression of a common core set of hypoxia-responsive genes underlies the improvement of aerobic performance across all life stages. Rather, it suggests that either different sets of genes underlie life-stage specific responses, or that a core set of genes is only transiently differentially expressed. We cannot distinguish these alternatives with our data, as it would require sampling individuals at different developmental stages and we only measured gene expression in adults. Nonetheless, our results demonstrate that exposure to hypoxia during prenatal development seems to blunt the transcriptional response to hypoxia during adulthood. This pattern is most evident in comparisons between the lifelong hypoxia and control groups where we found the fewest differentially expressed genes (Figure 1). In contrast, mice exposed only during adulthood exhibited a robust transcriptomic response, with over 6000 genes being differentially expressed in comparisons with the control mice. This pattern may indicate active remodeling of the diaphragm in mice exposed to hypoxia as adults, a process that may have been completed at earlier developmental timepoints in mice that were exposed as juveniles.

Many genes differentiate control mice from mice exposed to hypoxia at different developmental stages. In the lifelong hypoxia group, the top 10 differentially expressed genes include those involved in angiogenesis and axon guidance (Nrp1; Neuropilin 1) (Kofler & Simons, 2015), transmembrane ion transport (Slc26a3; Solute Carrier Family 26 Member 3), antigen binding/immune response (Ighv1; Immunoglobulin Heavy Variable 1), regulation of systemic osmotic pressure (Trpv4; Transient Receptor Potential Cation Channel Subfamily V Member 4) (Liedtke & Friedman, 2003), and steroid metabolism (Cyp3a13; Cytochrome P450 Family 3 Subfamily A Member 13) (Choudhary et al., 2003). Four enriched functional categories were Reactome categories related to antigen activity and response to tissue inflammation, which suggests a role in immunity.

In the post-natal hypoxia group, genes that were highly differentially expressed in comparison to the control group included functions such as regulation of cellular cholesterol metabolism (Pip4p1; Phosphatidylinositol-4,5-Bisphosphate 4-Phosphatase 1) (Hammond & Burke, 2020), calcium ion binding activity (Megf6; Multiple Epidermal Growth Factor-Like Domains Protein 6) (Teerlink et al., 2021), regulation of cell growth in differentiating tissues (Mxd4; MAX Dimerization Protein 4) (Boros et al., 2011), and axon guidance during neuronal development (Sema3f; Semaphorin 3F). Interestingly, Slc26a3, Cyp3a13, Megf6, and Trpv4 were all among the top most differentially-expressed genes for both lifelong and post-natal hypoxia exposures. Overall, the genes differentially-expressed in the post-natal hypoxia group involved immunity and Major Histocompatibility Complex functions, anatomical structure morphogenesis, and cellular response to oxygen-containing compounds (Table S4). In the adult hypoxia group, top differentially-expressed genes included some of those mentioned above (Pip4p1, Sema3f, Mxd4), as well as others related to myelin regulation and development of the diaphragm (Myrf; Myelin Regulatory Factor) (Rossetti et al., 2019).

### How are gene regulatory phenotypes associated with respiratory traits?

We identified several regulatory modules in the diaphragm that were associated with functional variation and improvements in arterial oxygen saturation under hypoxia. Of the five regulatory modules we identified, the expression of three was significantly associated with arterial oxygen saturation (M1, M2, M3) or ventilatory traits (M1, M2, M4), and the expression of one (M4) was significantly associated with relative diaphragm mass. Combined, the modules associated with improved oxygen saturation contained a total of 13,161 genes and were enriched for genes that participate in several relevant functions, including those related to responses to chemical stimulus and vasodilation (M1), mitochondrial function and organization (M2), and gene regulation and metabolic processes (M3) (Table S6). Similarly, module M4, which was significantly associated with relative diaphragm mass, tidal volume, and total ventilation, was enriched for genes that regulate transcriptional processes that could contribute to muscle growth. Interestingly, these modules are associated with physiological variation (e.g., arterial O_2_ saturation) across all three hypoxia treatments, and may represent common responses to hypoxia across life stages. However, previous experiments in deer mice have shown that distinct physiological mechanisms underlie improvements in aerobic performance resulting from developmental hypoxia (Ivy et al., 2021), but we did not observe any significant module correlations between gene expression and VO_2_max. This may have arisen because improvements in aerobic performance resulted from the small number of differentially expressed genes that were specific to developmental hypoxia (Figure 1), and/or because important transcriptomic changes occurred at earlier developmental stages that were not assessed here.

Genes that are highly connected or centrally located within co-regulatory networks may have particular functional importance in regulating trait expression, because their expression is highly correlated with many others within the network.We identified hub genes within each of the trait-associated modules, and this analysis revealed that these highly connected, centrally-located genes participate in a variety of processes that could influence diaphragm form and function. For example, cyclin L2 (*Ccnl2*) is the hub gene for module M4, which is correlated with relative diaphragm mass. *Ccnl2* regulates apoptosis (Zhuo et al., 2009), and patterns of allele frequency variation are indicative of a history of positive selection in the high-elevation population (Schweizer et al., 2021, 2019). Similarly, the hub gene for module M1 is calsyntenin 2 (*Clstn2*) which plays a key role in synaptic assembly and transmission (Lipina et al., 2016). These and other highly connected genes are promising targets for follow-up study and manipulative experiments.

As outlined above, each of the trait-associated modules exhibited life-stage specific responses to hypoxia exposure. For the modules positively associated with trait values (M1 and M3; Figure 4), the greatest expression was found in mice exposed to hypoxia during adulthood, followed by the postnatal and life-long exposures, respectively (Figure 3). The opposite pattern was found for those modules that were negatively associated with trait values (M2 and M4); the lowest expression of these modules was found in mice exposed to hypoxia during adulthood, followed by postnatal and life-long exposures (Figure 3).

### Have genes associated with ventilatory traits contributed to a past response to altitude-related selection?

Hundreds of genes in the trait-associated modules bear signatures of positive selection in the highland population, suggesting that allelic variation at these loci may have contributed to hypoxia adaptation. Twenty-three of the 207 DE genes that were common to all three treatment groups were also candidate genes for high-altitude adaptation in Schweizer et al. (2021) (Table SX). These genes have a number of functions, such as ion transport, apoptosis, immune function, and nervous system development. For example, ENSPEMG00000019815 (orthologous to *Phospholipase C Like 2*) may be a negative regulator of cold-induced thermogenesis; in hibernating ground squirrels, PLCL2 is moderately expressed in brown adipose tissues (Hampton et al., 2013), and has been previously shown to be differentially expressed in locomotory muscle of adult deer mice exposed to hypoxia (Cheviron et al., 2014). Similarly, TCF4 (*Transcription factor 4*), which causes Pitt-Hopkins Syndrome in humans (Amiel et al., 2007) was differentially expressed in all three treatment groups relative to the control group. A major symptom of Pitt-Hopkins Syndrome is hyperventilation (Amiel et al., 2007), suggesting that variation at this locus may be important in ventilatory responses to hypoxia.

Of the set of 209 genes that were differentially expressed in the developmental exposures, but not in the adult exposure, 16 genes were also within the PBS outliers (Table S7), suggesting that they may play key roles in developmental processes that are adaptive under hypoxia. For example, one gene of interest is *Cdon* (Cell adhesion molecule-related/down-regulated by oncogenes), which encodes a cell adhesion molecule that is necessary for proper development of skeletal muscles (Cole et al., 2004; Takaesu et al., 2006). Another notable gene is *Foxred1* (FAD Dependent Oxidoreductase Domain Containing 1), which is necessary for assembly of NADH dehydrogenase in the mitochondrial respiratory chain. Loss-of-function mutations within *Foxred1* cause human Complex I deficiency, which presents in infancy or early adulthood (Calvo et al., 2010; Lemire, 2015).

We also found hundreds of genes involved in WGCNA regulatory modules that are candidates for high-altitude adaptation. These genes include *Ccln2* (hub gene for M4), which we highlighted above, plus a number of other genes that participate in relevant pathways. For example, the M1 module contained 662 genes that were also candidates for selection; the functions of some of these genes include the GO cateogry “ion transmembrane transporter activity” (Table S10). The M2 module, which contained 407 selection candidates, was enriched for several GO categories related to cellular metabolic processes and intracellular organelles. Additionally, no GO categories were enriched in the 150 selection candidates within M3 (Table S10), nor within the 63 genes from M4. It is feasible that these genes may be affecting function by influencing membrane physiology (e.g., ion transport in cell membrane or organelles of muscle or neurons) or energy metabolism.

### Conclusions

We identified a core set of 207 genes that were differentially expressed in response to hypoxia regardless of the developmental timing of exposure. However, the vast majority of expression changes were unique to a particular developmental timepoint. Similarly, the magnitude of the transcriptomic response to hypoxia depended on the developmental onset of exposure. The most pronounced transcriptional response was found in mice exposed to hypoxia during adulthood, while mice exposed starting at earlier developmental timepoints expressed regulatory phenotypes that were more similar to normoxia-exposed control mice. These plastic changes in gene expression were associated with increased arterial oxygen saturation, which is expected to improve aerobic performance under hypoxia, and with changes in breathing. Moreover, hundreds of genes in trait-associated modules bear signatures of positive selection in high-elevation deer mice, providing a suggestive line of indirect evidence that allelic variation in genes associated with beneficial physiological changes have contributed to hypoxia adaptation.

## Acknowledgements

The authors would like to thank Drs. J. Velotta, K. Wilsterman, E. Moore, E. Kopania, and members of the Cheviron lab for helpful feedback on data processing and analysis. This work was funded by the following grants: to CMI from the Natural Sciences and Engineering Research Council of Canada (NSERC) PGS and OGS; to JFS from the National Institutes of Health (R01 HL159061) and the National Science Foundation (IOS-2114465 and OIA-1736249); to GRS from the Natural Sciences and Engineering Research Council of Canada (NSERC) Discovery Grant (RGPIN-2018-05707) and Canada Research Chairs Program (950-232589); to ZAC from the National Science Foundation (OIA 1736249 and IOS-1755411) and the National Institutes of Health (R15 HD103925).

## Data Accessibility Statement

Raw RNAseq data in fastq format is available from the NCBI Short Read Archive (pending accession #SUB12098471). Gene count and phenotypic trait data and scripts for their analysis are available on github at https://github.com/renaschweizer/pman_diaphragm_rnaseq.

## Benefit-Sharing Statement

Benefits from this research include the sharing of our data and results on public databases as described above, as well as the development and inclusion of international collaborative partnerships.

## Author Contributions

R.M.S., G.R.S., J.F.S., and Z.A.C. designed the research in consultation with C.M.I. and C.N. C.M.I., C.N., and J.F.S. collected tissue samples. J.F.S. coordinated sequencing of transcriptomes. C.M.I. and G.R.S. provided phenotype data. R.M.S. analyzed the data and performed the research through discussion with Z.A.C. R.M.S. and Z.A.C. wrote the paper, and all authors provided feedback on and approved of subsequent versions.

## Supporting Information

**Table S1**. List of selection candidate genes from Schweizer et al. 2019.

**Table S2.** Sequencing metrics for 35 RNAseq libraries, including quality score, percent bases above Q30, %GC content, number of reads, genome alignment rate (%), and assignment rate (%).

**Table S3.** Differentially expressed genes for life-long hypoxia, post-natal hypoxia, and adult hypoxia relative to control. For each gene, the Ensembl ID (Gene), log FC (log 2 fold change relative to control), log CPM (log 2 counts per million), FDR (False Discovery Rate-adjusted p-value), *Mus* Ensembl ID, and *Mus* gene name is provided.

**Table S4**. Results of *gProfiler* gene ontology enrichment analysis on lifelong hypoxia, post-natal hypoxia, adult hypoxia, only developmental hypoxia (lifelong and post-natal), and intersection gene lists.

**Table S5.** Pearson’s correlation and FDR-corrected p-values for association of WGCNA modules with diaphragm mass, arterial O_2_ saturation, and ventilatory phenotypes.

**Table S6**. Results of *gProfiler* gene ontology enrichment analysis on gene lists in four modules from WGCNA that had significant treatment effects.

**Table S7**. Differentially expressed genes in “developmental only” treatments (lifelong hypoxia and post-natal hypoxia) that overlap with selection candidate genes from Schweizer et al. 2019.

**Table S8**. Differentially expressed genes in intersection of three treatments (lifelong hypoxia, post-natal hypoxia, and adult hypoxia) that overlap with selection candidate genes from Schweizer et al. 2019.

**Table S9**. Sets of genes within four WGCNA regulatory modules that overlap with selection candidate genes from Schweizer et al. 2019.

**Table S10**. Results of *gProfiler* gene ontology enrichment analysis on gene lists in four modules from WGCNA that had significant treatment effects and that overlap with selection candidate genes from Schweizer et al. 2019.

**Figure S1.** Plot of scale independence and mean connectivity at soft threshold values of 1 to 30 as output from WGCNA analysis. We chose a soft thresholding power of 13, since this was the initial inflection point at which the scale free topology model fit began to decrease with higher thresholding powers

