## Supplementary figures and images for "Gene regulatory changes underlie developmental plasticity in respiration and aerobic performance in highland deer mice"

### Figure S1

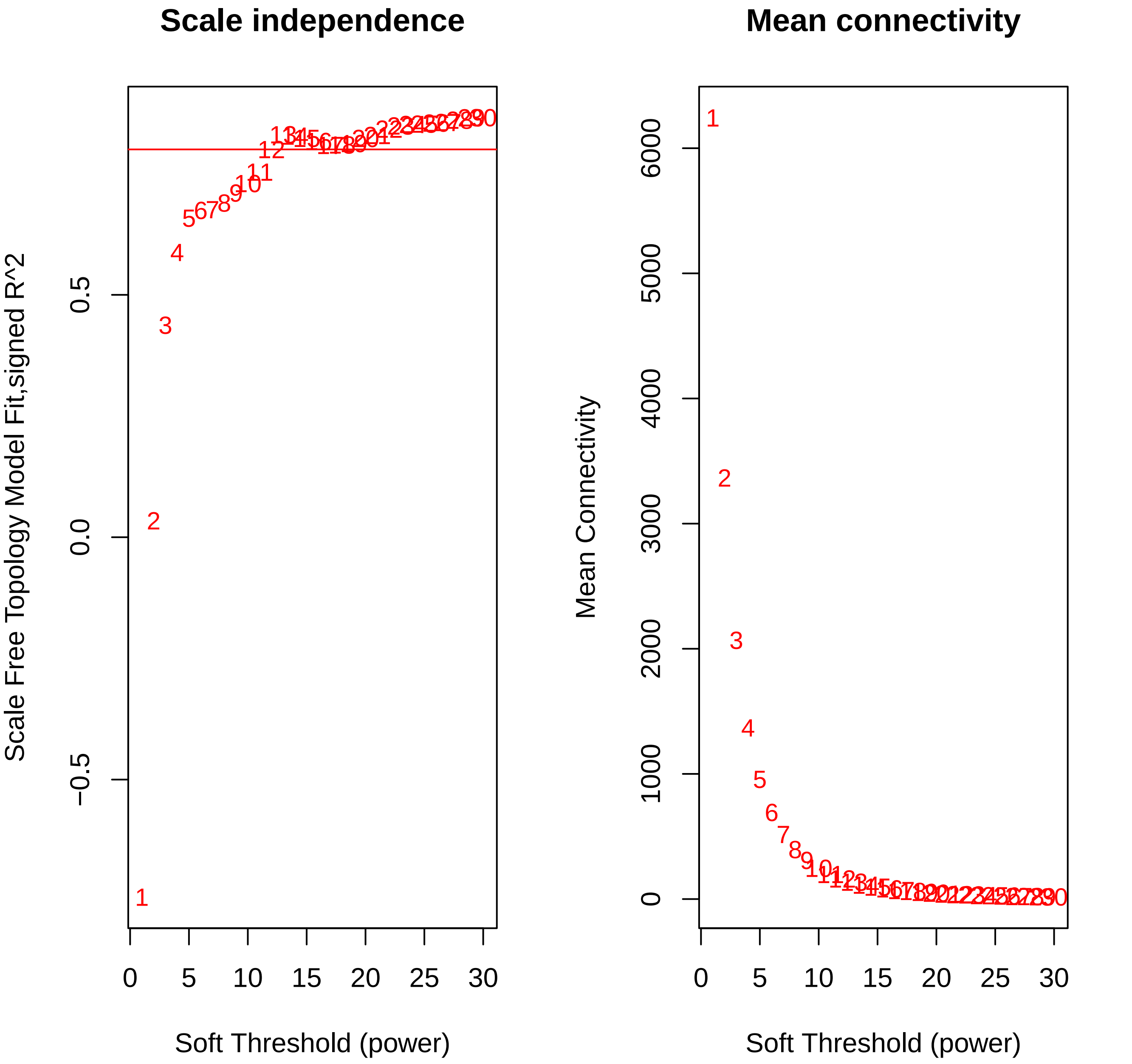
